# How to feed a little sparrow? Disentangling determinants and consequences of feeder use in the Eurasian tree sparrow *Passer montanus* (Linnaeus, 1758) using RFID technology

**DOI:** 10.64898/2025.12.14.694133

**Authors:** Adrián Sánchez Albert, Gabriel Munar Delgado, Francisco Pulido

## Abstract

There is extensive debate on supplemental bird feeding given the mixed evidence about its effects on avian taxa. In this study, we quantified feeder use in the Eurasian Tree Sparrow (*Passer montanus*) from a population in Central Spain using radio frequency identification (RFID) with the aim of unravelling its determinants and consequences. We found extensive variation in feeder use, both individually and temporally. Individuals that made use of feeders later spent more time feeding. Feeder use did not explain variation in reproductive success in our population. Our study indicates that inter-individual differences combined with seasonality are the main determinants of feeder use.

## 1. Introduction

Food availability is widely recognised as one of the fundamental factors limiting bird populations (Newton, 1980, 1998; Martin, 1987; White, 2008), further affecting reproductive success and survival of many bird species (Lack 1947; Cody, 1966; Martin, 1987; Grames *et al*., 2023). One important and easily accessible sources of food are humans and their settlements, from where many species obtain food either eventually or intentionally (Orams, 2002; reviewed by Oro *et al*., 2013; Becker *et al*., 2015).

Intentional supplementary bird feeding by humans (hereafter bird feeding; Jones, 2017; Reynolds *et al*., 2017) is a complex and controversial phenomenon, only recently subjected to academic and societal scrutiny (Jones & Reynolds, 2008; Robb *et al*., 2008a; Shutt & Lees, 2021). Several case-studies, reviews and meta-analyses were systematically conducted to dimension the impacts of bird feeding on bird species (*e*.*g*., Schoech & Hahn, 2007; Robb *et al*., 2008a; Ruffino *et al*., 2014; Murray *et al*., 2016; Reynolds *et al*., 2017; Francis *et al*., 2018; Plummer *et al*., 2019; Shutt & Lees, 2021; Grames *et al*., 2023) and proved somewhat contradictory (Robb *et al*., 2008a; Ruffino *et al*., 2014). While most studies have reported benefits (enhancement of reproductive output, *e*.*g*., Robb *et al*., 2008b; increased individuals health, Knutie, 2020), others have found no clear fitness benefits (*e*.*g*., Sonnenberg *et al*., 2023), or described negative direct effects on fitness (*e*.*g*., Chamberlain *et al*., 2008), and a plethora of other undesired indirect effects (*e*.*g*. habituation and possible feeder-dependency, increase in disease transmission, hyperpredation; Reynolds *et al*., 2017; Shutt & Lees, 2021). The reasons for this inconsistency among studies are not well known, but as suggested by Ruffino *et al*. (2014) they may relate to variation in 1) the life-history of the species (*e*.*g*., inadequate food for targeted species), 2) the environmental conditions (*e*.*g*., sufficient naturally available food) or 3) the experimental design (*e*.*g*., lack of statistical power).

The joint use of Passive Integrated Transponders (PIT, Gibbons & Andrews, 2004) and Radio-Frequency Identification (RFID, see Bridge & Bonter, 2011), as tagging and data-recording methods respectively, has a number of advantages for studying of the implications of bird feeding (Wilmers *et al*., 2015; Katzner & Arletazz, 2020; Lahoz-Monfort & Magrath, 2021). This technology is not only harmless for birds (*e*.*g*., great tits, Nicolaus *et al*., 2008; house sparrows, Schroeder *et al*., 2008), and is increasingly accessible for ornithologists (Bridge & Bonter, 2011), but it is also particularly well suited to study the effects of bird feeding due to the possibility of automatic recording of individual feeder use without interfering with the birds (Bridge & Bonter, 2011), which allows a continuous recording for sufficient periods of time. With all this, RFID technology can provide an especially valuable opportunity to answer the question that is haunting ornithologists for some time now: Is bird feeding really beneficial for bird species or is it more intricate than that?

The Eurasian tree sparrow (Passer montanus Linnaeus, 1758; hereafter tree sparrow) is a small, cavity-nesting (Snow & Perrins, 1998) and sedentary (Deckert, 1962; Field & Anderson, 2004) songbird, that readily uses both artificial nesting sites (Cordero, 1993; García-Navas *et al*., 2008a, 2008b; Węgrzynowicz, 2012; von Post & Smith, 2015) and feeders installed on its territories (Tryjanowski *et al*., 2016; Fülöp *et al*., 2019, 2022a; Plummer *et al*., 2019), and therefore, it is an ideal study species for the study of bird feeding. This species is frequently classified as omnivorous (Cramp & Brooks, 1994; Snow & Perrins, 1998; García-Navas, 2016), as it feeds mostly on insects and arthropods during the reproductive period, but gradually switches to a seed-based diet outside the breeding season (Sánchez-Aguado, 1986; Veiga, 1990; Holland *et al*., 2006). Although the most basic aspects of its diet and foraging ecology are well known (Samtchuk, 1981; Summers-Smith, 1995), and research interest on this topic persists to these days (*e*.*g*., Barta *et al*., 2004; Mónus & Barta, 2010, 2011; Fülöp *et al*., 2019, 2022a), to date, no study has analysed ‘feeder use and its determinants or its potential effects on reproductive success (*e*.*g*., Field & Anderson, 2004; Siriwardena *et al*., 2007).

In this paper we present, to our knowledge, the first study on the use of bird feeders by tree sparrows, using RFID technology. Although, since Bonter & Bridge’s first review of RFID use in ornithological research (2011), where they reported a total of 7 published papers implementing this technology in bird feeders, the number of studies on birds making use of this technology has steadily increased in the last decade (*e*.*g*., Bonter *et al*., 2013; Bandivadekar *et al*., 2018; Siekiera *et al*., 2019; Lajoei *et al*., 2019; see Reynolds *et al*., 2017).

We had three main objectives: first, determining variation in the temporal use of feeders on an intermediate (monthly) and long (between pre-reproductive and reproductive periods) timescale. Second, we aimed at identifying the factors that may affect their use (*e*.*g*., distance and first visit to feeders), and finally, determining the effects that the use of bird feeders may have on the reproductive output.

## 2. Materials & Methods

### 2.1 Study area and nest-box arrangement

Fieldwork was conducted in El Encinar de San Pedro of la Casa de Campo (West of Madrid, Central Spain, 40°25’33.6”N, -3°45’8.99”W), a restricted-access area within the Casa de Campo public park (∼1,700 ha, 650 m asl; Figure 1.). The site is located within a stand of evergreen forest, primarily occupied by *Quercus ilex rotundifolia*, with an open understory of broom (*Retama sphaerocarpa*). This suburban holm oak woodland is characterised by the prevalence of hole-nesting species such as tits, tree-creepers and tree sparrows, with the latter being the most common (Alonso & Purroy, 1979; Sánchez & Tellería, 1988).

**Figure 1.**
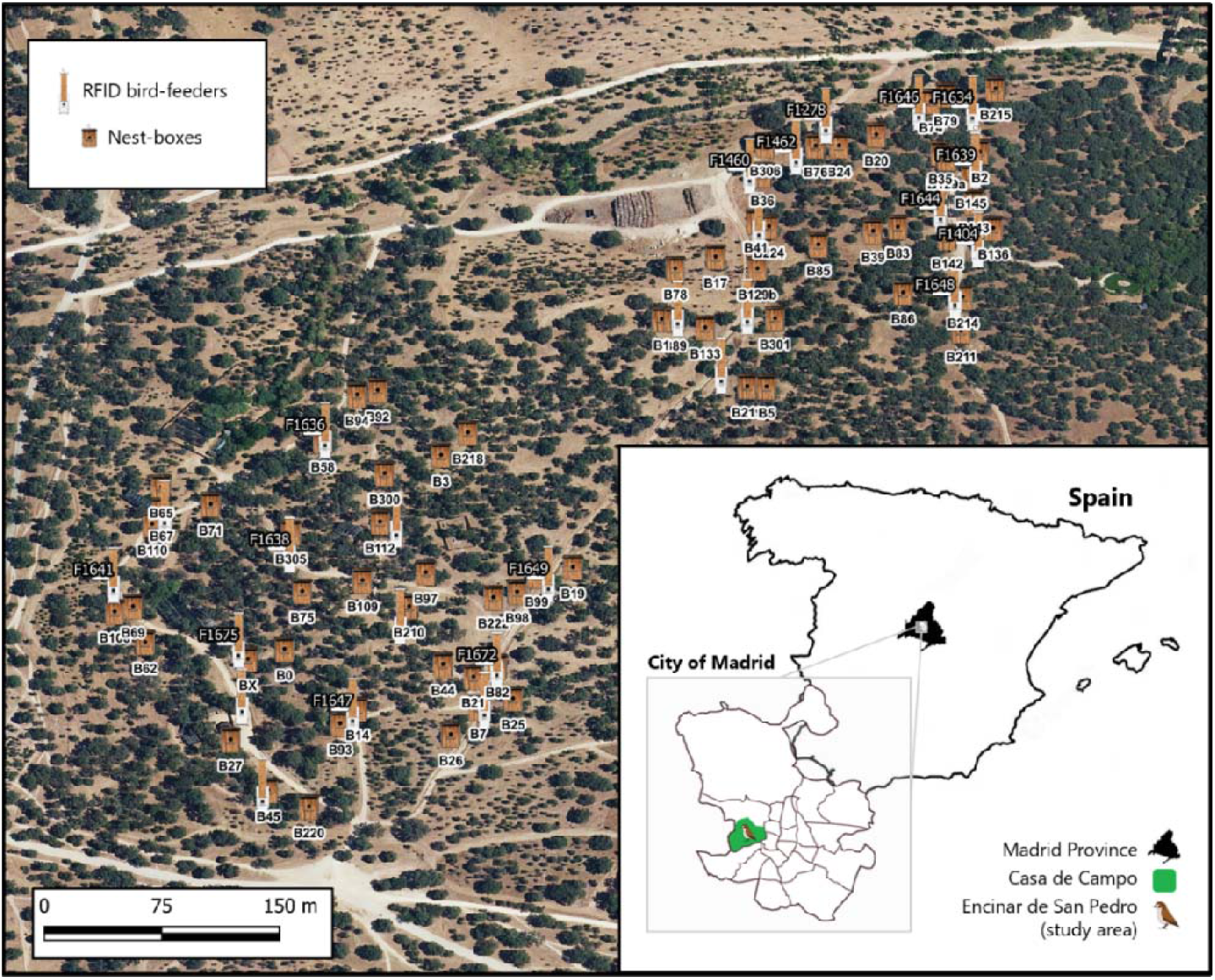
*Left*. Distribution of the 68 nest-boxes (BXXX) and the 26 RFID bird feeders (FXXXX) in the study area. *Bottom right*. Schematic map of the situation of the study area in the City of Madrid, the Autonomous Community of Madrid and Spain.

A total of 68 wooden nest boxes were installed in the study area in October 2019 (Figure 1). Nest-boxes were checked weekly from the end of March to mid-August 2021-22 to record the number of active nests, and data on reproduction (*e*.*g*., laying date, clutch size, fledging success). The study area was divided into two zones (‘North’ and ‘South’, 33 and 35 nest-boxes, respectively), separated by less than 1 km. This distribution of nest boxes was designed to test the Matching Habitat Choice Theory (MHC; Edelaar *et al*., 2008; 2017) for tree sparrows (Munar-Delgado, 2023).

### 2.2 RFID-based monitoring technology

Our monitoring technology system was composed of a set of readers (antennae/lectors), some integrated into automated bird feeders (26 in total, 13 in each zone), others removable for being relocated between nest-boxes (24 in total, 12 in each zone), and a set of external PIT-tags.

As of January 2020, individuals were captured either in nest boxes during night-roost inspections (outside the breeding period) or by mist-netting throughout the year. Upon capture, each bird was fitted with a single PIT-tag in a plastic leg ring (2 × 12 mm, circa 0.1 g, Figure 2c), which had a unique alphanumeric identification code. Therefore, every time a PIT-tagged bird passed through a data-logger in a nest-box or landed on a bird feeder, its identity, and date and time (with an accuracy of seconds) of each visit was detected by the RFID readers.

**Figure 2.**
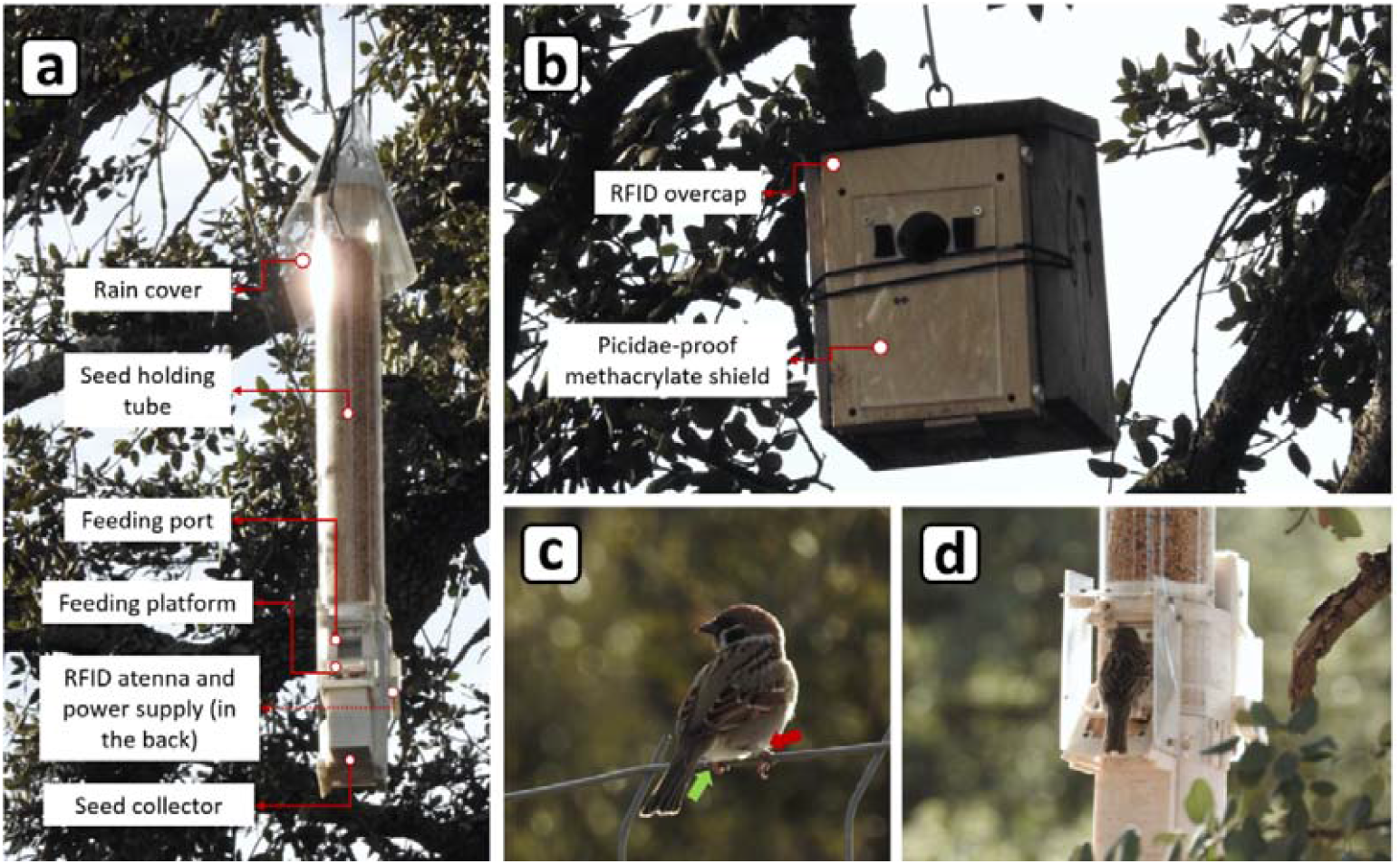
Examples of automated bird feeder (a) and nest-box (b) used in this study. Eurasian tree sparrows standing on a fence with an aluminium leg band (red arrow) in the right leg and a light green PIT-tag (hidden, green arrow) on the left leg (c), and feeding on a bird feeder (d). Photo credit: A. Sánchez-Albert.

Bird feeders were activated on 26th August, and their position was fixed on 16th September 2021. Feeders were designed to be closed with a lift-door by default, and to open automatically when the reader registered a specified PIT-tag code that granted access (set prior to the experiment). This design prevents that more than one bird could use the feeder at a time. Feeders were filled periodically with a commercial seed mix, so that food was continuously available. No bird feeders were present prior to the study, and, to our knowledge, no other sources of supplemental food were available in the study area and its surroundings.

All readers were checked weekly for data retrieval and battery replacement, and reconditioning/substitution if necessary. In addition, as the number of installed nest boxes exceeded the number of available readers (68 vs. 24), they were relocated between boxes (always within the same area) during the weekly visits, so that each breeding bird could be identified.

### 2.3 Data selection and variables

For this study we only considered data on the first breeding pairs (i.e., first mating) of the birds nesting in the installed nest-boxes, in order to discard uncontrolled variables related to mating dynamics (*e*.*g*., fitness, divorces, polygamy, etc.), as this species is multi-brooded (García-Navas & Sanz, 2012, García-Navas, 2016), polygamous (Cordero *et al*., 2002; Seress *et al*., 2007), and in which divorces are frequent (Sasvári & Hegyi, 2000; Arbelo-Cenalmor *et al*., 2022).

Birds were monitored during two consecutive years: in 2021 and 2022, and since one of the pairs was present in both years, our final sample size is 66 individuals, 33 males and 33 females (i.e., 33 breeding pairs). For their first breeding attempt, pairs occupied 11 (15.9%) and 23 (33.3%) out of the 69 installed nest-boxes in 2021 and 2022, respectively. Although they were not the only nesting species (some Paridae nests were found during nest-box monitoring), it was the prevalent species occupying nest-boxes during the study. In 2021, the first laying was observed in April (08/04), and in 2022 at the end of March (25/03). Accordingly, we defined the period January-March as the pre-reproductive (preR) period and April-June as the reproductive period (R). Sexing of individuals was possible through RFID identification during nocturnal incubation, as this is done exclusive by females (García-Navas & Sanz, 2012).

Our variables of feeder use were computed out of RFID detection data for each month and PIT-tagged bird using Python v. 3.10.5 (Python Software Foundation, 2022) as programming language and Sublime Text 3 (Sublime HQ Pty Ltd, 2019) as code editor. These feeder variables were: (1) the number of different feeders visited (*N*_*feeders*_), (2) the number of visitats made to all visited feeders (*N*_*visits*_), (3) the time in seconds spent feeding from those (*T*_*seconds*_), and 4) the number of agonistic interactions in which the birds were involved during feeder use (*N*_*displacements*_), which did not allow differentiate between individuals attacking and attacked. Also, three derived variables were calculated from the previous variables: (5) the rate of displacements (*disp_rate = N*_*displacements*_ */N*_*visits*_), (6) the time spent feeding during each visit (*time_per_visit = T*_*seconds*_ */N*_*visits*_), and (7) and on each feeder (*time_per_feeder = T*_*seconds*_ */N*_*feeders*_). All these variables were anlayzed for differences by sex and month, and by periods (pre-reproductive, preR and reproductive, R). To account for the period changes in feeder use, these variables were calculated for both periods, and to study their effect on reproductive success they were compared by pairs, in those cases where both partners had a known PIT-tag. From all these variables, the time spent at the bird feeders was assumed to be the variable most accurately reflecting feeder use, given the foraging strategy of the tree sparrow, in which a single feeder visit implies the consumption of several seeds (i.e., “sit-and-eat” strategy, Barta *et al*., 2004; Mónus & Barta, 2011; sensu Hughes *et al*., 2021).

In addition to these seven variables related to feeder use, the distance between feeders and the breeding nest-boxes in meters were calculated with the function ‘Distance to nearest hub (points)’ using Quantum GIS software v.30 (QGIS, 2023), in order to calculate a mean distance to visited bird feeders (mean_*distance* = Σ(*D*_*feeders*_)/*N*_*feeders*_), during the two periods considered (preR and R).

Monitoring nest-boxes during the reproductive period allowed us to obtain six direct measures of reproductive success for each breeding pair: (1) number of clutches (*N*_*clutches*_), (2) number of eggs per clutch (*N*_*eggs/clutch*_), (3) number of chicks per brood (*N*_*chicks/brood*_), (4) the average weight of each brood (mean_*weight* = Σ(*W*_*chicks*_)/*N*_*chicks/brood*_), as well as the (5) total number of eggs layed (Σ(*N*_*eggs/clutch*_)), and (6) the total number of chicks hatched (Σ(*N*_*chicks/brood*_)). Also, we determined phenological variables related to the reproductive success (laying dates from the 1^st^ and 2^nd^ clutches), the time between clutches (*e*.*g*., days between 1^st^ and 2^nd^ breeding attempt), and the first visit made by each individual to a bird feeder.

### 2.4 Statistical analyses

#### 2.4.1. Temporal analysis

Due to our data’s temporal structure, independent (*e*.*g*., differences between years) and paired (*e*.*g*., differences between periods, months and sexes) sample tests were the prevalent analyses. Before conducting the analyses, normality and homoscedasticity of the variables were tested using Shapiro-Wilk’s and Levene’s test, respectively, in order to apply parametric or non-parametric statistics when appropriate.

In the case of independent sample tests (factor: year), fulfilment (or not) of normality and homoscedasticity was addressed by a manual stepwise statistic selection method: 1) fulfilment of both assumptions involved the use of Student’s t-test, 2) non-normality only implied the application of Mann-Whitney’s U, and 3) violation of both assumptions entailed the use of Welch’s T (Ruxton, 2006). On the other hand, in the paired samples analysis, a similar statistical selection method was employed, whereby: 1) normality was addressed with paired Student’s t-test, and 2) violation of this assumption with the Wilcoxon signed-rank test for two paired samples (Derryberry *et al*., 2017). Furthermore, when it was necessary to analyse the association between variables using correlation methods, given the systematic violation of normality, the rank-based alternative of Spearman’s Rho (ρ), was used (Puth *et al*., 2015). For conducting all these analyses, the solid version (v. 2.3.21) of statistical software Jamovi (Jamovi Project, 2022) was used.

#### 2.4.2. Analyses of the effects on feeder use and reproductive success

We conducted a manual modelling approach, fitting generalized linear models (GLMs) through the ‘glm’ function from the ‘stats’ package, in R version 4.3.0 (R Core Team, 2023), to analyse: (1) how potential predictor variables (*e*.*g*., distance, experience, date of first visit to feeders) were linked to feeder use (as time spent in feeders, T_seconds_), and (2) how variables related to feeder use could have influenced different measures of reproductive success (*e*.*g* number of eggs and chicks, mean brood weight, number of clutches, etc.) of our population. Since differences between periods and years were found (Supp. Table 5), we control them by including predictors matching the period of the dependent variable or by including ‘year’ as a factor.

For the analysis of the effects of explanatory variables on feeder use, 3 different models were fitted: one for the time spent at the feeders in the pre-reproductive period (GLM_PreR Time_), another for the reproductive period (Time-GLM_R Time_), and one for both periods together (Time-GLM_Total_ Time). Thus, we investigated whether effects of the explanatory variables differed between periods (reproductive and non-reproductive), as well as making a general prediction. In the case of feeder-related variables effects on reproductive output, 6 different GLMs were fitted, one for each variable reflecting reproductive success. We discerned between effects on reproductive success of the first clutch, i.e., clutch size (GLM_1st-Clutch Size_), brood size (GLM_1st-Brood Size_) and average weight of the brood (GLM_1st-Mean Weight_); and effects on the overall reproductive success of each pair, i.e., the total number of clutches (GLM_Total Clutches_) and the total amount of eggs laid (GLM_Total Eggs._) by stable pairs. In addition, we fitted another GLM for studying the possible effects of feeder use on the time interval between the laying of the first and second clutch (GLM_1st-2nd_). The error structure and link function used in each model is specified in Supplemental Tables 7 and 8.

Instead of transforming variables that were not normally distributed and fitting linear models (LMs), we implemented GLMs given the balance between flexibility/simplicity that they offer in their application (Pekár & Brabec, 2016, 2017), avoiding the difficulty to interpret transformed data (*e*.*g*., St-Pierre *et al*., 2018 for count data). Also, given the presence of missing values in our dataset, the Akaike Information Criterion (AIC) statistic could not be used to rank the degree of support for different regression models and thus for model selection (Nakagawa & Freckleton, 2011). For this reason, only one model was fitted for each dependent variable. However, the AIC-value for each model and the statistics of their respective null models are given in the summary table for each model (see Supp. Tables 7. and 8.).

## 3. Results

Of the 633 individuals ringed and tagged since January 2019, we registered 68 birds (34 pairs) laying their first clutch in the nest-boxes during our study (2021-2022). Among these breeding birds, only two individuals, one female (2021) and one male (2022) could not be identified, thus making *n*_*2021*_ = 21 and *n*_*2022*_ = 45 PIT-tagged breeding birds. These tagged birds made a total of 125,326 visits and spent 4,515,217 secsonds (∼1,250.23 hrs.) on bird feeders during the January-June period. In total bird feeders registered 184,089 visits, and 7,059,438 seconds (∼1,960.86 hrs.) of use. Thus, post-reproductive/autumn use (from August to December) accounted for 36.04% of the total (=2,544,221 seconds) and 31.92% of the feeding events (=58,763 visits).

All variables reflecting feeder use correlated positively with each other (see, Supp. Table 1.). The strongest correlation was between the number of visits and time spent on frequented feeders (ρ = 0.86, p < 0.001). Significant, but moderate correlations (ρ < 0.7) were found among the rest of the variables (see, Supp. Table 1.).

### 3.1. Individual variation in feeder use

Of all birds breeding in 2021, only one PIT-tagged birds (a female) was not detected by the RFID data loggers installed in the feeders, thus making n_2021_ = 20 individuals recorded at some point of the year using bird feeders. In 2022, 4 females and 2 males did not visit the feeders (remaining n2022 = 39).

We found substantial variability in monthly and daily visits among individual tree sparrows. Throughout the 12-month period, the average time birds spent at the feeders was 5,533 seconds (SD = 7,365.11, range = 0–69,603), and the mean number of visits per feeder was 154 (SD = 285, range = 0–5,779) per month. Birds made an average of 5.12 visits/day (SD = 5.98, range = 0–29.06 visits/day) and spent on average 184.45 seconds/day (SD = 207.16, range = 0– 967.39 seconds/day) on feeders. This variation was also reflected in the number of visited bird feeders, with a mean of 1.65 devices (SD = 1.15, range = 0-12) per individual and month.

Differences between sexes in feeder use were found to be mostly non-significant (see, Supp. Table 2.). Only the time spent at each visit in the pre-reproductive period was significantly different between sexes (W = 13, p = 0.005), with males spending more time at each feeding event before reproduction. Other variables such as the number of visits during the reproductive period (W = 184, p = 0.086) and the time spent at a feeder during the pre-reproductive period (F = -1.847, df = 14, p = 0.086) were close to significance.

### 3.2. Temporal variation in feeder use

#### 3.2.1. Variation in feeder use between periods and months

Significant differences in feeder use were detected when comparing the pre-reproductive period and the reproductive period: visiting frequency was higher (W = 486.5, p = 0.004, MD = -402.5), while aggressive interactions (measured both as frequency, W = 461, p < 0.001, MD = 136.68, and as a rate, W = 418, p < 0.001, MD = 18.31), and mean distance to bird feeders (F = 2.962, df = 44, p = 0.005, MD = 28.15), were lower during the reproductive period (see, Supp. Table 3.).

When we controlled for the sex, females showed a steadier use of feeders than males between periods, since only the number of feeders used by females differed between these two periods, being higher in the reproductive period (F = -2.134, df = 32, p = 0.041, MD = -1.242). Males, on the other hand, mirrored the general results described in the previous paragraph. They differed in the frequency of visits (W = 81, p = 0.01, MD = -700.42), agonistic interactions (frequency, W = 225, p < 0.001, MD = 138.5; rate, W = 208, p = 0.033, MD = 0.19), and mean distance to feeders (F = 2.573, df = 22, p = 0.017, MD = 30.33), and in the length of each visit, which was shorter in the reproductive period (W = 208, p = 0.033, MD = 24.2).

There were significant overall differences in monthly feeder use in both years, measured in both time spent feeding in bird feeders (Friedman’s Test, 2021: χ2 = 80.9, df = 5, p < 0.001; 2022, χ2 = 43.4, df = 5, p < 0.001), and the total number of visits made to feeders (Friedman’s ANOVA, 2021 χ2 = 112, df = 5, p < 0.001; 2022, χ2 = 46.9, df = 5, p < 0.001). The detailed pairwise results of this analysis (Durbin-Conover pair comparison) can be found in Supp. Table 4.

In 2021, despite the slight increase in feeder use within the pre-reproductive period (Figure 3a.), it remained more or less similar between months as no significant monthly differences in feeder use were found, when measured as frequency of visit and time spent (although between January-March was marginally significant, p = 0.085). In the reproductive period, by contrast, monthly usage was found to vary significantly between months, both in visits and time (Supp. Table 4., Tables 1A and 1B), increasing sharply until reaching a maximum in June (Figure 3a.). On the other hand, in 2022, the pre-reproductive period did show differences between January-February (time, p = 0.064; visits, p = 0.013), but not between January-March or February-March. In the reproductive period, usage also varied on a monthly basis (both measures tested, *i*.*e*., time and visits; see Supp. Table 4., Tables 2A and 2B) but in a different way than in 2021. In 2022, instead of increasing from March and reaching its maximum in June, it fell from March-April, reached a lower maximum in May and fell again in June (see, Figure 3b.).

**Figure 3.**
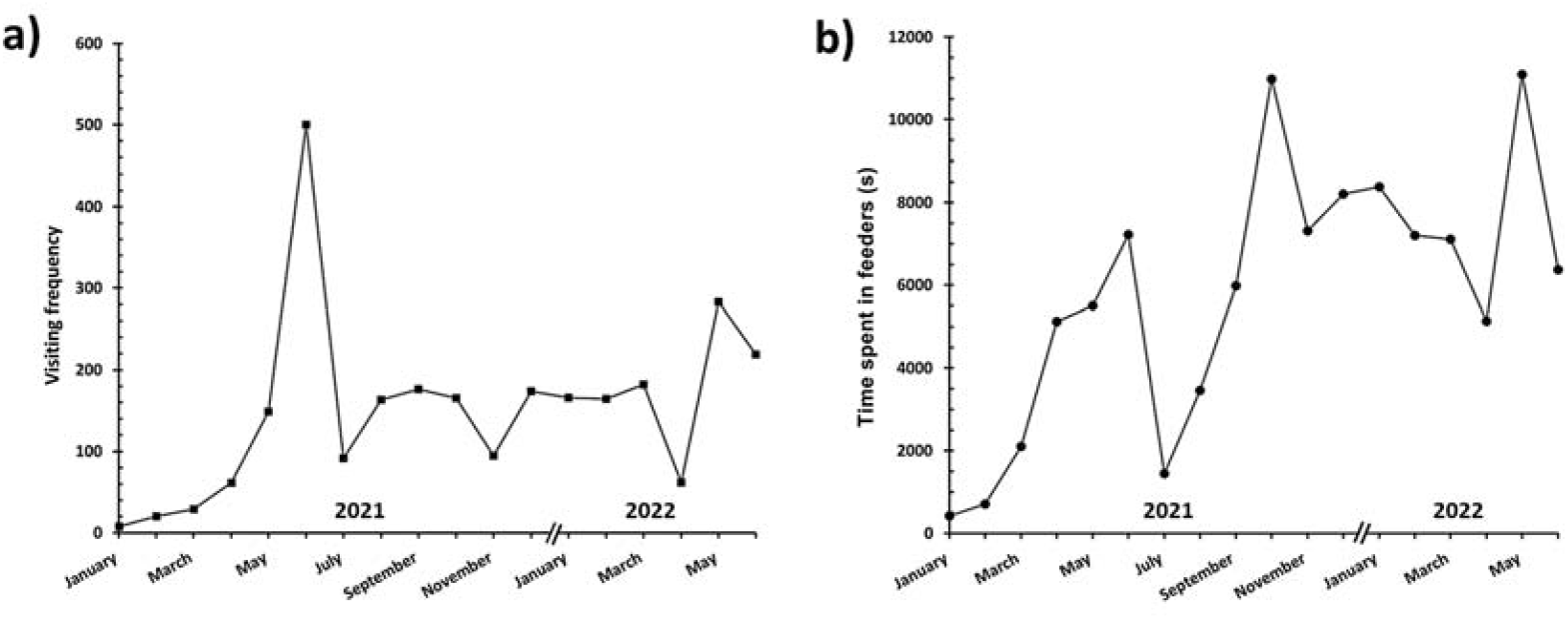
Monthly variation in mean feeder use represented by the time spent in feeders (in seconds, a) and number of visits (b) over months for which we have records.

#### 3.2.2. Among-year differences in feeder use and fitness

Feeder use differed markedly, and in a complex manner, between the two years considered in this study. Each variable related to feeder use varied in a different way, yet total feeder use did not vary between 2021 and 2022, except for the number of feeders visited each year (see, Supp. Table 5.). To illustrate this complexity, we can have a closer look at two different measures of feeder use, *i*.*e*., the number of visits made and the number of feeders visited. While in the pre-reproductive period of 2021 birds made fewer visits to the feeders than in 2022 (F = -4.11, df = 55.7, *p* < 0.001, *MD* = -509.75), in the reproductive period the opposite is true (F = -2.63, df = 23.23, *p* = 0.015, *MD* = 1144.43). A different result was found for the number of feeders visited; it was higher in 2022, overall and in both periods (see, Supp. Table 5.).

There were no among-year differences in clutch and brood size, and mean brood weight of the first breeding attempt. Total reproductive success, *i*.*e*., the number of clutches made, number of laid eggs and raised chicks per pair, was higher in 2022. Despite this, the first visit of birds to feeders was significantly earlier in 2022 than in 2021 (F = 4.32, df = 56.8, p < 0.001, MD = 143).

### 3.3. Relationship between feeder use and potential determinants

#### 3.3.1. Correlates of feeder use: distance and agonistic interactions

In the pre-reproductive period, time spent per feeder correlated negatively with mean distance (*ρ* = -0.33, *p* = 0.015, see Figure 4.a). However, in the reproductive period, although also negative, the correlation between these variables was found to be non-significant (*ρ* = -0.16, *p* = 0.24, Figure 4.b.). In this same period, time spent per visit correlated positively with mean distance to the visited feeders (*ρ* = 0.278, *p* = 0.017), but not in the pre-reproductive period, in which the correlation resulted slightly negative and non-significant (*ρ* = -0.152, *p* = 0.323). A correlation was also found between mean distance in both periods, with those individuals that used more distant bird feeders in the pre-reproductive period also using distant feeders in the reproductive period (*ρ* = 0.678, *p* < 0.001) (see, Supp. Table 6A.).

**Figure 4.**
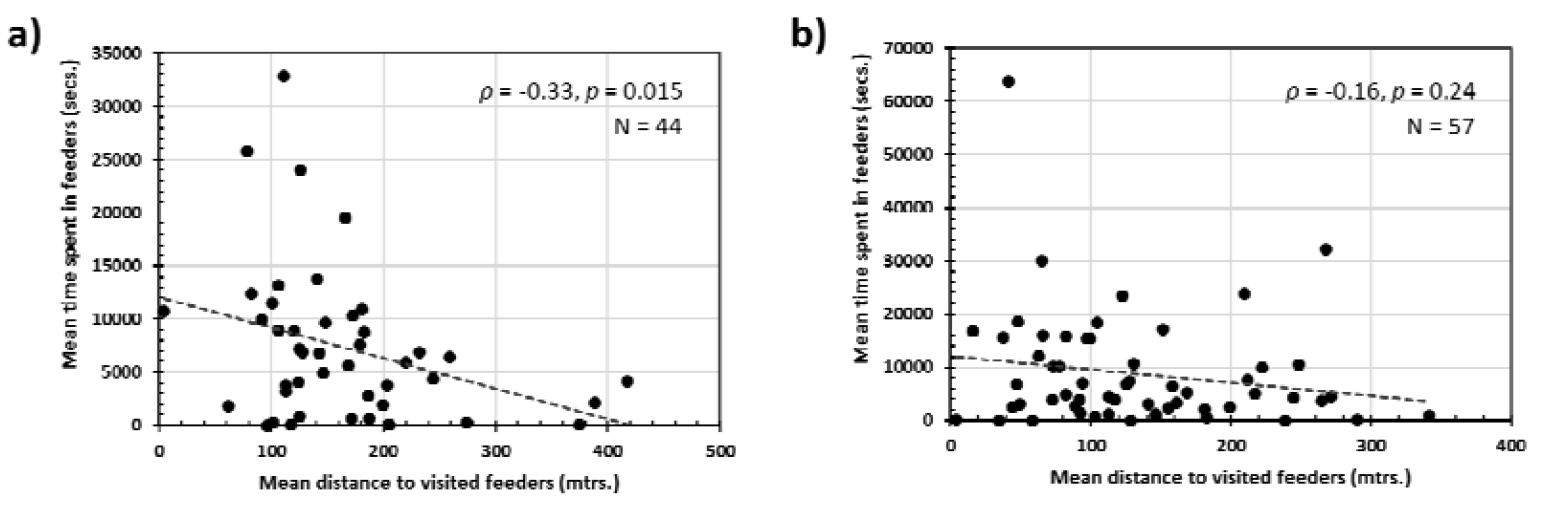
Relationship between mean distance and mean time spent at bird feeders by individuals, both in the pre-reproductive (a) and reproductive (b) periods. Direction (+/-) and strength of the correlation (Spearman’s rho, *ρ*), p-value and sample size of each analysis are shown.

Displacements (*i*.*e*., agonistic interactions; Supp. Table 6B.), generally, correlated positively with the number of feeders used (*preR, ρ* = 0.555, *p* < 0.001; *R, ρ* = 0.300, *p* = 0.022), the amount of time spent using them (*preR, ρ* = 0.464, *p* = 0.004; *R, ρ* = 0.462, *p* < 0.001) and the number of feeders visited in both periods (*preR, ρ* = 0.504, *p* = 0.001; *R, ρ* = 0.523, *p* < 0.001). In the reproductive period, this was also true for the mean time spent on visited feeders (*ρ* = 0.297, *p* = 0.024). Also, there was a strong positive correlation for displacements between periods (*ρ* = 0.77, *p* < 0.001), suggesting that those individuals involved in more agonistic interactions in the pre-reproductive period were also those involved in more interactions during the reproductive period. Furthermore, there was a negative correlation between the number of interactions and the mean distance between breeding nest-box and feeders visited, but this relation was only significant in the reproductive period (*preR, ρ* = -0.226, *p* = 0.215; *R, ρ* = -0.278, *p* = 0.032).

#### 3.3.2. Generalized linear models of feeder use

Total time spent on bird feeders (GLMTotal Time, df = 41, missing values = 26) was positively associated with the factor year, although marginally (*p* = 0.068), while the other considered predictors, including distance and date of first visit, showed non-significant associations with this measure of feeder use. When time spent in feeders was restricted to the pre-reproductive period (GLMPreR Time, df = 41, mv = 26), sex (*p* = 0.031) and first date of visit (*p* = 0.012) turned out to be significant predictors, whereby males spent less time on the feeders, and a later first use meant a greater amount of time spent feeding on the supplemental resource. Unlike the general model, mean distance to the bird feeders used in *preR* was marginally associated (*p* = 0.092) with the time of feeder use, year being non-significant (p = 0.275). During the reproductive period the total time of feeder use (GLMR Time, df = 52, mv = 15) was predicted by year (p = 0.037) and marginally by the first visit (p = 0.082). See Supp. Table 7. for a detailed summary on the statistics of each GLM.

### 3.4. Relationship between feeder use and fitness variables

For the variables reflecting reproductive success of the first breeding event, none of the variables related to the use of feeders included in the models were significant (see, the summary tables of the models GLM_1st-Clutch Size_, GLM_1st-Brood Size_ and GLM_1st-Mean Weight_ in Supp. Table 8.). A similar conclusion can be drawn for those models aiming at predicting overall reproductive success fitting means of feeder use (see, GLM_Total Eggs_, GLM_Total Clutches_) and models predicting the time interval between the first and second clutch (GLM_1st-2nd_, see Supp. Table 8).

## 4. Discussion

Our study revealed that tree sparrows showed a wide variation in their use of bird feeders, with large inter-individual and temporal differences, both at long (years) and intermediate (periods/months) timescales. We then found some additional factors that explained, to some extent this variation, like the date of first visit to feeders, the mean distance to feeders, etc., but we found no significant effects of feeder use on reproductive success.

### 4.1. Inter-individual and temporal variation drive feeder use

Large inter-individual differences found on feeder use is likely to be a reflection of the covariation between personality and foraging behaviour (Sih *et al*., 2012; Toscano *et al*., 2016), which has already been found for the tree sparrow during feeding in flocks (Fülöp *et al*., 2019). This variation is congruent with similar studies of other bird species (*e*.*g*., Poecille atricapillus, Latimer *et al*., 2018; Lajoei *et al*., 2019, and Baelophus bicolor, Bonter *et al*., 2013; Parus major and Cyanistes caeruleus, Crates *et al*., 2016), although tree sparrows made fewer daily visits to visited feeders. This lower feeder use compared to other bird species may be due to different foraging strategies: passerids (Family Passeridae, *e*.*g*., tree sparrow) spend some time consuming several seeds on the feeder, and parids (Fam. Paridae) take less time as they pick up a seeds for later consumption, and therefore they are expected to make more visits (“sit and eat” vs. “grab and go” strategies, sensu Hughes *et al*., 2021). Another explanation may be the availability of sufficient natural food in our study area, in contrast to other latitudes where, during winter, snow can severely limit access to natural seed banks (*e*.*g*., Croston *et al*., 2016).

In relation to sex differences, males and females only differed in the time spent per visit during the pre-reproductive period, probably because males gain preferential access to resources through intersexual aggressions (Lee & Sung, 2023). The lack of marked differences in feeder use, especially during the breeding period, is in line with the synchronicity of the pairs, whose members equally share the tasks of parental care (Sasvári & Hegyi, 1994; García-Navas *et al*., 2016).

The temporal feeders use dynamics by tagged tree sparrows, both monthly and between periods, were consistent with the annual life cycle of the species. Regarding our findings on changes in mean distance to feeders, a previous study in Japan (Sano, 1973) also reported a decrease in foraging distance during the breeding season, when chick provisioning becomes a priority. Although distance to visited feeders may seem a poor estimate of general foraging range, a recent study has demonstrated that the movements of GPS-tagged tree sparrows are highly conditioned by the position of bird feeders (Fülöp *et al*., 2022b). With respect to agonistic interactions for feeder access, although tree sparrows often attack or threaten conspecifics (Torda *et al*., 2004; Mónus & Barta, 2010), in some cases with fatal outcomes (*e*.*g*., Lee *et al*., 2022), this behaviour mainly occurs during the pre-breeding period. Therefore, the decrease of these interactions that we detected is congruent with what would be expected: it takes place when the dominance hierarchy in the population is already established (Torda *et al*., 2004; Mónus *et al*., 2016; Lee & Sung, 2023), and tree sparrows become strictly solitary (Mónus & Barta, 2010). Finally, the counterintuitive increase in feeder use during the reproductive period could be explained by the ability of parents to modulate their foraging activity (Williams, 2018): individuals would adjust their investment in self-feeding seeds from feeders and searching for insects to feed the chicks by visiting closer feeders.

#### 4.2. The role of feeder distance

We found that feeder use by breeding tree sparrows, in addition to being determined by individual behaviour and life cycle, it was also related to other predictors. Feeder use, expressed as the time spent on each feeder, showed a negative association with the mean distance to the feeders, suggesting that foraging behaviour of tree sparrows is conditioned by perceived risk of predation, which increases with distance from shelter (Tsurim, 2005; Tsurim *et al*., 2008), and could be magnified by the role that bird feeders sometimes play as ecological traps, attracting predators (*e*.*g*., Malpass *et al*., 2016; Hanmer *et al*., 2016). However, in our population, this factor did not prove to be particularly limiting, and this may be due to the fact that the distances between nest-boxes (i.e., shelters) and the nearest feeders were relatively small, when compared to the species home range (Field & Anderson, 2004).

#### 4.3. Fitness consequences of feeder use

Despite the pronounced variation in feeder use by tree sparrows, we did not find any evidence for an effect of feeder use on reproductive success, as previous case-studies report for other bird species (*e*.*g*., Cyanistes caeruleus and Parus major, Crates *et al*., 2016; Poecile gambei, Sonnenberg *et al*., 2023). Nevertheless, we suggest that bird feeding could have a masked effect on our target population, favouring poor-quality individuals to the point of reaching reproductive outputs typical of high-quality individuals (see, Crates *et al*., 2016). This hypothesis is supported by the fact that tree sparrows delayed in using feeders, and probably arriving late at this area in winter, which are widely thought to be subordinated to older conspecifics and relegated to sub-optimal territories (Pinowski *et al*., 2008), spent more time feeding on this supplemental resource.

#### 4.4. Limitations of the study and future directions

Our bird feeder network equipped with RFID technology allowed us to study the use of feeders by tree sparrows on a fine scale (Bonter & Bridge, 2011), with an accuracy of seconds. However, our feeder network suffered exceptional limitations during the pre-reproductive period of 2021, caused by the combined effect of extreme weather events (i.e., January, 2021, Storm Filomena and the consequent cold wave, Smart, 2021) and COVID-19 restrictions (Aubry *et al*., 2021). These events made it impossible to access the study area, and hence, to repair and/or replace those feeders directly damaged by the snowstorm (*e*.*g*., freezing, fallen from tree branches, etc.), and refilling their seed containers. These technical constrains may have caused individuals to stop recognising feeders as a predictable food source, and hence to drastically reduce their reliance on this resource, especially to those whose seed reserves were totally depleted, reflecting to some extent the results of Mady *et al*. (2021).

A lower daily frequency of visits to bird feeders, when compared with other species data, may raise concern about the detection probability of our RFID systems (Hughes *et al*., 2021). However, detection performance was tested by cross-checking with video recordings, and resulted highly correlated with observed visits (>90%, n = 15 RFID antennas; Cabezas-Serrano *et al*., 2023), in congruence with other RFID systems (*e*.*g*., >99%, Firth *et al*., 2018; 100%, Falk *et al*., 2021). However, given that tree sparrows are less likely to make quick visits to the feeder (3 s on average; Hughes *et al*., 2021), and that our feeders automatically restrict access to food depending on bird identification, we would expect the detection capability of our technology to be a priori high.

To mitigate potential shortcomings of this study, future studies should, (1) explore the relationship between feeder use and its predictors and consequences through inferentially robust methodologies (*e*.*g*., BACI analysis, Fenn *et al*., 2020; longitudinal experiments, Sonnenberg *et al*., 2023); and (2) study the effects of different, and probably more suitable, supplementary food types (Ruffino *et al*., 2014). Although bird feeders usually provide seeds that may favour adult tree sparrows during winter (Siriwardena *et al*., 2007; Xingjun *et al*., 2018), provisioning of protein-rich food (*e*.*g*., animal prey), should be a more adequate measure for increasing fledgling success and chick condition (*e*.*g*., Passer domesticus, Peach *et al*., 2014), and potentially help to mitigate the slow but steady decline of their populations (Ramos-Elvira *et al*., 2023).

## 5. Conclusion

This study demonstrates, on one hand, the virtues of RFID technology, which allowed us to constantly measure feeder use by tree sparrows and explain its variation through individual identity, reproductive state, date of first visit, etc., and on the other hand, the limitations of our methodology, which cannot help resolving the question of whether bird feeding is inherently beneficial to our study species, highlighting the need for applying experimental methodology. Nevertheless, we proved that bird feeding has no detrimental effects on our study population, and reinforce the idea that casuistry (cf. Ruffino *et al*., 2014; Shutt & Less, 20221) governs discussion on the effects of bird feeding on bird populations.

## Acknowledgments

Special thanks to Alvar, Jessica Arbelo Cenalmor, Laura Cabezas Serrano and Javier who thoroughly worked on the maintenance of the feeder and nestbox network, and helped in making field work less harsh.

