## Supplementary Material for "How to feed a little sparrow? Disentangling determinants and consequences of feeder use in the Eurasian tree sparrow *Passer montanus* (Linnaeus, 1758) using RFID technology"

**Supplemental Material**

**Supplemental Table 1.-** Correlation matrix between measures of overall feeder use (number of visits, agonistic interactions, time spent and number of feeders visited). Results of the normality tests for each variable (Shapiro-Wilk test) are shown, as well as the correlation coefficient (Spearman's rho) and the p-value are shown.

|  | Normality Test | | Statistics | *visits_tot* | *disp_tot* | *time_tot* | *n_feeder_tot* |
| --- | --- | --- | --- | --- | --- | --- | --- |
|  | W | *p* |  |  |  |  |  |
| *visits_tot* | 0.855 | < .001 | Rho (ρ) | — |  |  |  |
|  |  |  | *p-value* |  |  |  |  |
| *disp_tot* | 0.704 | < .001 | Rho (ρ) | 0.589 | — |  |  |
|  |  |  | *p-value* | < .001 |  |  |  |
| *time_tot* | 0.893 | < .001 | Rho (ρ) | 0.857 | 0.611 | — |  |
|  |  |  | *p-value* | < .001 | < .001 |  |  |
| *n_feeder_tot* | 0.95 | 0.01 | Rho (ρ) | 0.518 | 0.567 | 0.519 | — |
|  |  |  | *p-value* | < .001 | < .001 | < .001 |  |

**Supplemental Table 2.-** Summary table of the differences on feeder use between sexes for which paired two-sample tests were used. The statistic used depended on the result of the normality test (Shapiro-Wilk test) on each case: non-normal variable, Wilcoxon signed-rank test; normal variable, Student's t-test (*). The difference of means and their confidence intervals are provided for all analysis, and degrees of freedom (df) are provided for those t-tests. Note: low *p-values* in the normality test reflect the violation of the assumption.

|  | Paired two-sample test | | | | | Normality test | |
| --- | --- | --- | --- | --- | --- | --- | --- |
|  | Statistic | *p-value* | ES, Mean difference (Balance: female - male) | Confidence interval | | W | *p-value* |
|  |  |  |  | Lower | Upper |  |  |
| *visits_female - male_preR* | -0.156* (df = 30) | 0.877 | -20.5806 (−) | -289.809 | 248.648 | 0.934 | 0.056 |
| *visits_female - male_R* | 184 | 0.086 | -300.5 (↑) | -847 | 52.5 | 0.888 | 0.003 |
| *disp_female - male_preR* | -0.736* (df = 15) | 0.473 | -39.3125 (−) | -153.231 | 74.606 | 0.944 | 0.399 |
| *disp_female - male_R* | 90 | 0.385 | -10.50004 (−) | -34 | 16 | 0.919 | 0.048 |
| *n_female - male_feeder_preR* | -1.056* (df = 31) | 0.299 | -0.875 (−) | -2.565 | 0.815 | 0.936 | 0.058 |
| *n_female - male_feeder_R* | 0.454* (df = 31) | 0.653 | 0.3125 (−) | -1.093 | 1.718 | 0.973 | 0.586 |
| *time_female - male_preR* | -1.847* (df = 14) | 0.086 | -19430.0667 (↑) | -41991.811 | 3131.678 | 0.955 | 0.604 |
| *time_female - male_R* | 0.952* (df = 24) | 0.351 | 9171.52 (−) | -10709.24 | 29052.28 | 0.921 | 0.055 |
| *time_female - male_per_visit_preR* | 13 | 0.005 | -32.56109 (↑) | -64.4563 | -8.4907 | 0.755 | 0.001 |
| *time_female - male_ per_visit_R* | 0.843* (df = 25) | 0.407 | 6.803 (−) | -9.813 | 23.419 | 0.948 | 0.205 |
| *disp_female - male_ratio_preR* | 36 | 0.182 | -0.1554 (−) | -0.4857 | 0.3736 | 0.698 | < .001 |
| *disp_female - male_ ratio_R* | 106 | 0.516 | -0.00779 (−) | -0.0431 | 0.0249 | 0.276 | < .001 |
| *time_female - male_per_feeder_preR* | -1.715* (df = 14) | 0.108 | -3982.8947 (−) | -8964.881 | 999.091 | 0.914 | 0.155 |
| *time_female - male_ per_feeder_R* | 146 | 0.672 | -894.96667 (−) | -4331.3417 | 4619.0429 | 0.839 | 0.001 |
| *mean_female - male_distance_feeders_preR* | 0.725* (df = 14) | 0.481 | 8.6663 (−) | -16.987 | 34.319 | 0.941 | 0.397 |
| *mean_female - male_distance_feeders_R* | -0.41* (df = 24) | 0.686 | -8.8221 (−) | -53.27 | 35.626 | 0.926 | 0.071 |
| *date_female - male _first_feeder_visit* | -0.24* (df = 25) | 0.813 | -9.6154 (−) | -92.281 | 73.05 | 0.98 | 0.881 |

**Supplemental Table 3.-** Summary table of the differences in feeder use between periods (pre-reproductive, *preR* and reproductive, *R*) for which paired two-sample tests were used. For details of the statistics used, see Appendix 2.

|  | Paired two-sample test | | | | | Normality test | |
| --- | --- | --- | --- | --- | --- | --- | --- |
|  | Statistic | *p-value* | Mean difference (*MD*, Balance: preR -R) | Confidence interval | | W | *p-value* |
|  |  |  |  | Lower | Upper |  |  |
| Individuals | | | | | | | |
| *visits_preR - R* | 486.5 | 0.004 | -420.5 (↑) | -780.5001 | -121.5 | 0.824 | < .001 |
| *disp_preR - R* | 461 | < .001 | 138.6787 (↓) | 77 | 210 | 0.89 | 0.003 |
| *n_feeder_preR - R* | 580.5 | 0.584 | -0.4999 (−) | -1.5001 | 1 | 0.941 | 0.003 |
| *time_preR - R* | -1.305* (df = 43) | 0.199 | -8394.205 (↑) | -21366.5616 | 4578.1525 | 0.965 | 0.193 |
| *time_per_visit_preR - R* | 544 | 0.575 | 2.8985 (↓) | -7.3757 | 21.023 | 0.737 | < .001 |
| *disp_ratio_preR - R* | 418 | < .001 | 18.3145 (↓) | 11.8521 | 32.867 | 0.555 | < .001 |
| *time_per_feeder_preR - R* | 352 | 0.097 | -2465.0272 (↑) | -5248.3838 | 327.569 | 0.887 | < .001 |
| *mean_distance_feeder_preR - R* | 2.962* (df = 44) | 0.005 | 28.145 (↓) | 8.9981 | 47.2914 | 0.954 | 0.072 |
| By sex: female | | | | | | | |
| *visits_female_preR - R* | 169 | 0.195 | -226 (↑) | -677 | 113.5 | 0.933 | 0.042 |
| *disp_female_preR* | 108 | 0.142 | 65 (↓) | -23.5 | 195.5 | 0.806 | < .001 |
| *n_female_feeder_preR - R* | -2.134* (df = 32) | 0.041 | -1.242 (↑) | -2.4285 | -0.0563 | 0.95 | 0.134 |
| *time_female_preR - R* | 65 | 0.082 | -25866 (↑) | -44675.5 | 2154 | 0.901 | 0.037 |
| *time_female_per_visit_preR - R* | 66 | 0.089 | -10.3734 (↑) | -22.5912 | 4.319 | 0.887 | 0.019 |
| *disp_female_ratio_preR – R* | 115 | 0.433 | 0.0235 (−) | -0.0245 | 0.183 | 0.436 | < .001 |
| *time_female_per_feeder_preR - R* | 68 | 0.103 | -3732.2667 (↑) | -10793.1667 | 439.75 | 0.821 | 0.001 |
| *mean_female_distance_feeders_preR - R* | 1.709* (df = 20) | 0.103 | 27.3 (↓) | -6.0287 | 60.6293 | 0.931 | 0.144 |
| By sex: male | | | | | | | |
| *visits_male_preR - R* | 81 | 0.01 | -700.418 (↑) | -1597 | -130 | 0.772 | < .001 |
| *disp_male_preR* | 225 | < .001 | 138.4999 (↓) | 73 | 206 | 0.783 | < .001 |
| *n_male_feeder_preR - R* | 164.5 | 0.221 | 1 (−) | -1 | 2.5 | 0.869 | < .001 |
| *time_male_preR – R* | 0.8*(df = 22) | 0.432 | 6223.739 (↓) | -10764 | 21588.5 | 0.953 | 0.338 |
| *time_male_per_visit_preR - R* | 208 | 0.033 | 24.2049 (↓) | 1.3332 | 70.501 | 0.746 | < .001 |
| *disp_male_ratio_preR – R* | 225 | < .001 | 0.1913 (↓) | 0.1185 | 0.49 | 0.624 | < .001 |
| *time_male_per_feeder_preR - R* | 117 | 0.54 | -1464.722 (−) | -5060.3889 | 2840.537 | 0.905 | 0.032 |
| *mean_male_distance_feeders_preR - R* | 2.573* (df = 22) | 0.017 | 30.332 (↓) | 7.3882 | 53.529 | 0.963 | 0.521 |

**Supplemental Table 4.** Monthly differences on feeder use between years (2021 = 1, 2022 = 2) and measured as time in seconds (A), and the number of visit (B) Paired tests were conducted using the non-parametric Durbin-Conover pair comparison test.

| 1.A | Statistic | *p-value* | 1.B | Statistic | *p-value* |
| --- | --- | --- | --- | --- | --- |
| *January - February* | 0.241 | 0.81 | *January - February* | 0.124 | 0.902 |
| *January - March* | 1.729 | 0.085 | *January - March* | 1.072 | 0.284 |
| *January - April* | 3.297 | 0.001 | *January - April* | 2.268 | 0.024 |
| *January - May* | 5.508 | < .001 | *January - May* | 5.36 | < .001 |
| *January - June* | 8.282 | < .001 | *January - June* | 10.473 | < .001 |
| *February - March* | 1.488 | 0.138 | *February - March* | 0.948 | 0.344 |
| *February - April* | 3.055 | 0.002 | *February - April* | 2.144 | 0.033 |
| *February - May* | 5.267 | < .001 | *February - May* | 5.236 | < .001 |
| *February - June* | 8.041 | < .001 | *February - June* | 10.349 | < .001 |
| *March - April* | 1.568 | 0.118 | *March - April* | 1.196 | 0.233 |
| *March - April* | 3.779 | < .001 | *March - April* | 4.288 | < .001 |
| *March - June* | 6.553 | < .001 | *March - June* | 9.401 | < .001 |
| *April - May* | 2.211 | 0.028 | *April - May* | 3.092 | 0.002 |
| *April - June* | 4.985 | < .001 | *April - June* | 8.205 | < .001 |
| *May - June* | 2.774 | 0.006 | *May - June* | 5.113 | < .001 |
| 2.A | Statistic | *p-value* | 2.B | Statistic | *p-value* |
| *January - February* | 1.857 | 0.064 | *January - February* | 2.511 | 0.013 |
| *January - March* | 1.2969 | 0.196 | *January - March* | 1.861 | 0.064 |
| *January - April* | 4.1856 | < .001 | *January - April* | 4.343 | < .001 |
| *January - May* | 2.5349 | 0.012 | *January - May* | 2.363 | 0.019 |
| *January - June* | 1.2085 | 0.228 | *January - June* | 0.384 | 0.701 |
| *February - March* | 0.56 | 0.576 | *February - March* | 0.65 | 0.516 |
| *February - April* | 2.3286 | 0.02 | *February - April* | 1.832 | 0.068 |
| *February - May* | 4.3919 | < .001 | *February - May* | 4.874 | < .001 |
| *February - June* | 0.6485 | 0.517 | *February - June* | 2.127 | 0.034 |
| *March - April* | 2.8886 | 0.004 | *March - April* | 2.482 | 0.014 |
| *March - April* | 3.8319 | < .001 | *March - April* | 4.225 | < .001 |
| *March - June* | 0.0884 | 0.93 | *March - June* | 1.477 | 0.141 |
| *April - May* | 6.7205 | < .001 | *April - May* | 6.706 | < .001 |
| *April - June* | 2.9771 | 0.003 | *April - June* | 3.959 | < .001 |
| *May - June* | 3.7434 | < .001 | *May - June* | 2.747 | 0.006 |

**Supplemental Table 5.**  Differences in the feeder use among-years. The table also reports the normality (Shapiro-Wilk test) and homoscedasticity (Levene’s test) tests, and the pertinent statistics based on the violation of these assumptions (see, *2.4.1. Temporal analysis*).

|  | Shapiro-Wilk Test | | Levene’s Test | | | | T-Test for independent samples: Mann-Whitney’s U (^u^), Welch’s T (^a^) or Student’s T (*) | | | | | |
| --- | --- | --- | --- | --- | --- | --- | --- | --- | --- | --- | --- | --- |
|  | W | *p-value* | F | df | df2 | *p* | Statistic | df | *p-value* | ES, Mean difference (Balance: 2021 - 2022) | Confidence interval | |
|  |  |  |  |  |  |  |  |  |  |  | Lower | Upper |
| Bird feeder related variables (individuals) | | | | | | | | | | | | |
| *visits_preR* | 0.83 | < .001 | 14.2783 | 1 | 64 | < .001 | -4.1084^a^ | 55.71 | < .001 | -509.7524 | -758.335 | -261.1702 |
| *visits_R* | 0.87 | < .001 | 19.2274 | 1 | 64 | < .001 | 2.628^a^ | 23.23 | 0.015 | 1144.4317 | 244.065 | 2044.7983 |
| *visits_tot* | 0.901 | < .001 | 4.9869 | 1 | 64 | 0.029 | 1.2868^a^ | 28.44 | 0.209 | 634.6794 | -374.94 | 1644.2992 |
| *disp_preR* | 0.869 | < .001 | 0.7091 | 1 | 33 | 0.406 | 68.5^u^ | - | 0.125 | -77.0001 | -222 | 67.99999 |
| *disp_R* | 0.808 | < .001 | 3.0178 | 1 | 44 | 0.089 | 176^u^ | - | 0.68 | -3 | -35.9999 | 27.99997 |
| *disp_tot* | 0.82 | < .001 | 0.3894 | 1 | 45 | 0.536 | 168^u^ | - | 0.311 | -52.4465 | -208 | 30.00005 |
| *time_preR* | 0.938 | 0.019 | 0.8077 | 1 | 42 | 0.374 | 170.5^u^ | - | 0.776 | -3271 | -22996 | 16660.99995 |
| *time_R* | 0.92 | < .001 | 10.0848 | 1 | 56 | 0.002 | 1.4217^a^ | 20.9 | 0.17 | 15922.2783 | -7374.694 | 39219.2507 |
| *time_tot* | 0.932 | 0.003 | 1.5798 | 1 | 56 | 0.214 | 318.5^u^ |  | 0.614 | 5986 | -22148 | 37850.00002 |
| *n_feeder_preR* | 0.951 | 0.011 | 14.5279 | 1 | 64 | < .001 | -4.6794^a^ | 63.08 | < .001 | -3.4254 | -4.888 | -1.9626 |
| *n_feeder_R* | 0.96 | 0.032 | 0.0272 | 1 | 64 | 0.87 | 344^u^ | - | 0.076 | -2 | -3.0001 | 4.36E-05 |
| *n_feeder_tot* | 0.975 | 0.198 | 0.5921 | 1 | 64 | 0.444 | -2.8192* | 64 | 0.006 | -2.9302 | -5.0065 | -0.85379 |
| *mean_distance_feeders_preR* | 0.911 | 0.002 | 2.0563 | 1 | 43 | 0.159 | 173^u^ | - | 0.725 | -8.2217 | -54.7201 | 35.74184 |
| *mean_distance_feeders_R* | 0.956 | 0.034 | 0.1222 | 1 | 57 | 0.728 | 304^u^ | - | 0.383 | -19.2024 | -62.8021 | 24.31376 |
| *mean_distance_feeders_tot* | 0.955 | 0.081 | 2.4633 | 1 | 43 | 0.124 | -0.7063* | 43 | 0.484 | -18.4911 | -71.2857 | 34.3036 |
| *date_first_feeder_visit_mod* | 0.951 | 0.02 | 4.77 | 1 | 57 | 0.033 | 4.32^a^ | 56.8 | < .001 | 143 | 76.5 | 209 |
| Reproductive success-related variables | | | | | | | | | | | | |
| *pair_tot_clutches* | 0.913 | 0.01 | 0.66093 | 1 | 32 | 0.422 | 72.5^u^ |  | 0.033 | -0.9999 | -1 | -4.92e−6 |
| *pair_st_clutch_size* | 0.652 | < .001 | 0.07244 | 1 | 32 | 0.79 | 123.5^u^ |  | 0.915 | 3.55E-05 | -4.03e−5 | 4.53E-05 |
| *pair_st_brood_size* | 0.795 | < .001 | 1.92785 | 1 | 32 | 0.175 | 116^u^ |  | 0.687 | -1.20e−5 | -1 | 1 |
| *pair_st_mean_clutch_weight* | 0.627 | < .001 | 1.91023 | 1 | 28 | 0.178 | 85^u^ |  | 0.906 | 0.0383 | -2.04 | 1.38 |
| *pair_egg_tot* | 0.948 | 0.108 | 0.79483 | 1 | 32 | 0.379 | -2.208* | 32 | 0.035 | -3.5415 | -6.809 | -0.2745 |
| *pair_chick_tot* | 0.954 | 0.189 | 0.00209 | 1 | 30 | 0.964 | -2.068* | 30 | 0.047 | -2.6818 | -5.33 | -0.0337 |
| *pair_mean_weight_tot* | 0.978 | 0.721 | 0.0634 | 1 | 31 | 0.803 | -1.3705* | 31 | 0.18 | -1.6327 | -4.0625 | 0.79697 |
| *pair_st_laying_date* | 0.411 | < .001 | 8.36 | 1 | 32 | 0.011 | 1.387^a^ | 10.12 | 0.195 | 50.5 | -30.4 | 131 |
| *pair_nd_laying_date* | 0.568 | < .001 | 39.8604 | 1 | 18 | < .001 | -0.98^a^ | 3 | 0.399 | 99.3 | -223.1 | 422 |
| *days_btw_st_nd_laying_dates* | 0.951 | 0.388 | 1.6813 | 1 | 18 | 0.211 | 0.8587* | 18 | 0.402 | 1.5 | -2.1698 | 5.16983 |

**Supplemental Table 6.** Correlation matrices between five measures of feeder use and (A) mean distances to visited feeders, and (B) the number of agonistic interactions during feeder visits. Both matrixes show feeder use split by periods and also report the correlation of the main variable between periods.

| 6A. Correlations between distance and feeder use | | | | | | | | | |
| --- | --- | --- | --- | --- | --- | --- | --- | --- | --- |
| Pre-reproductive period | | *visits* | *time_per_visit* | *time* | *time_per_feeder* | *n_feeder* | Comparison between periods | | *mean_distance_feeders_preR* |
| *mean_distance_feeders* | Rho (ρ) | 0.003 | -0.152 | -0.161 | -0.328 | 0.389 | *mean_distance_feeders_R* | Spearman’s Rho | 0.678 |
|  | *p-value* | 0.985 | 0.323 | 0.296 | 0.015 | 0.004 |  |  |  |
| Rreproductive period | | *visits* | *time_per_visit* | *time* | *time_per_feeder* | *n_feeder* |  |  |  |
| *mean_distance_feeders* | Rho (ρ) | -0.031 | 0.278 | 0.066 | -0.157 | 0.533 |  | *p-value* | < .001 |
|  | *p-value* | 0.408 | 0.017 | 0.688 | 0.243 | < .001 |  |  |  |

| 6B. Correlations between displacements and feeder use | | | | | | | | | | |
| --- | --- | --- | --- | --- | --- | --- | --- | --- | --- | --- |
| Pre-reproductive period | | *visits* | *time_per_visit* | *time* | *time_per_feeder* | *n_feeder* | *mean_distance_feeders* | Comparison between periods | | *disp_preR* |
| *disp* | Rho (ρ) | 0.504 | 0.089 | 0.464 | 0.162 | 0.555 | -0.226 | *disp_R* | Rho (ρ) | 0.767 |
|  | *p-value* | 0.001 | 0.628 | 0.004 | 0.375 | < .001 | 0.215 |  |  |  |
| Rreproductive period | | *visits* | *time_per_visit* | *time* | *time_per_feeder* | *n_feeder* | *mean_distance_feeders* |  |  |  |
| *disp* | Rho (ρ) | 0.523 | -0.135 | 0.462 | 0.297 | 0.300 | -0.278 |  | *p-value* | < .001 |
|  | *p-value* | < .001 | 0.376 | < .001 | 0.024 | 0.022 | 0.032 |  |  |  |

**Supplemental Table 7.** Fitted GLMs inferring feeder use predictors for three time periods (B. pre-reproductive, C. reproductive period and A. both) and their correspondent null models, with their associated AIC values and the difference in AIC (ΔAIC) between both. Significance, t-values and estimates are provided for each fitted GLM. Significant variables are shown in bold with an asterisk (*) and marginally significant variables are shown in bold with a dot (.).

| A. GLM_Total TIme_ | | | | | | | | | |
| --- | --- | --- | --- | --- | --- | --- | --- | --- | --- |
| Model no. | Explanatory Variables | K | ΔAIC | | df (obs. Deleted) | | Null Dev. | | AIC |
| Dependent variable: *time_tot* | | | | | | | | | |
| 1 | *null* | 0 | - | | 57 (10) | | 112.3 | | 1389 |
| 2 | ***year*** *(.) + sex (ns) + experience (ns) + date_first_feeder_visit (ns) + mean_distance_feeders_tot (ns)* | 5 | 365 | | 41 (26) | | 19.87 | | 1024 |
| Model summary | | | | | | | | | |
|  | | Estimate | | Std. error | | *t* value | | *P-*value | |
| Intercept | | -1.171e-02 | | 6.079e-03 | | -1.927 | | **0.0574 .** | |
| *year2022* | | 5.799e-06 | | 3.007e-06 | | 1.928 | | **0.0618 .** | |
| *montanus_sexmale* | | -2.438e-07 | | 2.389e-06 | | -0.102 | | 0.9193 | |
| *montanus_experienceyes* | | 1.894e-06 | | 3.072e-06 | | 0.617 | | 0.5414 | |
| *mean_distance_feeders_tot* | | 2.578e-08 | | 2.034e-08 | | 1.267 | | 0.2132 | |
| *date_first_feeder_visit* | | 1.436e-08 | | 1.127e-08 | | 1.274 | | 0.2109 | |

| B. Time-GLM_PreR Time_ | | | | | | | | | |
| --- | --- | --- | --- | --- | --- | --- | --- | --- | --- |
| Model no. | Explanatory Variables | K | ΔAIC | | df (obs. Deleted) | | Null Dev. | | AIC |
| Dependent variable: *time_preR* | | | | | | | | | |
| 1 | *null* | 0 | - | | 43 (24) | | 65.8 | | 1011 |
| 2 | *year (ns) +* ***sex*** *(*) + experience (ns) +* ***date_first_feeder_visit*** *(*) +* ***mean_distance_feeders_preR*** *(.)* | 5 | 43.8 | | 41 (26) | | 65.51 | | 967.2 |
| Model summary | | | | | | | | | |
|  | | Estimate | | Std. error | | *t* value | | *P-*value | |
| Intercept | | -2.432e-02 | | 2.196e-02 | | -1.107 | | **0.0346 *** | |
| *year2022* | | 1.205e-05 | | 1.087e-05 | | 1.109 | | 0.2750 | |
| *montanus_sexmale* | | -1.949e-05 | | 8.698e-06 | | -2.241 | | **0.0313 *** | |
| *montanus_experienceyes* | | 1.052e-05 | | 1.066e-05 | | 0.987 | | 0.3304 | |
| *mean_distance_feeders_preR* | | 1.053e-07 | | 6.088e-08 | | 1.730 | | **0.0922 .** | |
| *date_first_feeder_visit* | | 9.293e-08 | | 3.493e-08 | | 2.660 | | **0.0116 *** | |

| C. Time-GLM_R Time_ | | | | | | | | | |
| --- | --- | --- | --- | --- | --- | --- | --- | --- | --- |
| Dependent variable: *time_R* | | | | | | | | | |
| 1 | *null* | 0 | - | | 57 (10) | | 109.1 | | 1323 |
| 2 | ***year*** *(*) + sex (ns) + experience (ns) +* ***date_first_feeder_visit*** *(.) + mean_distance_feeders_R (ns)* | 5 | 104 | | 52 (15) | | 78.84 | | 1219 |
| Model summary | | | | | | | | | |
|  | | Estimate | | Std. error | | *t* value | | *P-*value | |
| Intercept | | -3.879e-02 | | 1.811e-02 | | -2.141 | | 0.2750 | |
| *year2022* | | 1.920e-05 | | 8.960e-06 | | 2.142 | | **0.0374 *** | |
| *montanus_sexmale* | | 8.223e-06 | | 7.730e-06 | | 1.064 | | 0.2929 | |
| *montanus_experienceyes* | | 4.976e-06 | | 8.977e-06 | | 0.554 | | 0.5820 | |
| *mean_distance_feeders_R* | | 5.529e-08 | | 5.628e-08 | | 0.982 | | 0.3310 | |
| *date_first_feeder_visit* | | 5.008e-08 | | 2.817e-08 | | 1.778 | | **0.0820 .** | |

**Supplemental Table 8.** GLMs inferring effects of feeder use on six measures of reproductive success and their correspondent null models. See Appendix 7, for table details.

| A. GLM_1st-Clutch Size_ | | | | | | | | | |
| --- | --- | --- | --- | --- | --- | --- | --- | --- | --- |
| Model no. | Explanatory Variables | K | ΔAIC | | df (obs. Deleted) | | Null Dev. | | AIC |
| Dependent variable: *1^st^_clutch_size* | | | | | | | | | |
| 1 | *null* | 0 | - | | 33 (0) | | 47.02 | | 49.02 |
| 2 | *time_female_preR (ns) + date_female_first_feeder_visit_mod (ns) + pair_st_laying_date_mod (ns) + pair_mean_distance_feeders_preR (ns)* | 4 | 23.08 | | 14 (19) | | 20.19 | | 25.94 |
| Model summary | | | | | | | | | |
|  | | Estimate | | Std. error | | *t* value | | *P-*value | |
| Intercept | | 9.348e+00 | | 5.752e+00 | | 1.625 | | 0.104 | |
| *time_female_preR* | | -5.115e-05 | | 3.637e-05 | | -1.406 | | 0.160 | |
| *date_female_first_feeder_visit_mod* | | -4.571e-03 | | 4.435e-03 | | -1.031 | | 0.303 | |
| *pair_st_laying_date_mod* | | -7.072e-02 | | 5.074e-02 | | -1.394 | | 0.163 | |
| *pair_mean_distance_feeders_preR* | | -3.340e-03 | | 4.922e-03 | | -0.679 | | 0.497 | |

| B. GLM_1st-Brood Size_ | | | | | | | | | |
| --- | --- | --- | --- | --- | --- | --- | --- | --- | --- |
| Model no. | Explanatory Variables | K | ΔAIC | | df (obs. Deleted) | | Null Dev. | | AIC |
| Dependent variable: *1^st^_brood_size* | | | | | | | | | |
| 1 | *null* | 0 | - | | 33 (0) | | 16.18 | | 131.8 |
| 2 | *pair_time_preR (ns) + pair_st_laying_date_mod (ns) + pair_mean_distance_feeders_preR (ns)* | 3 | 69.35 | | 14 (19) | | 3.274 | | 62.46 |
| Model summary | | | | | | | | | |
|  | | Estimate | | Std. error | | *t* value | | *P-*value | |
| Intercept | | 1.396e+00 | | 9.577e-01 | | 1.457 | | 0.145 | |
| *pair_time_preR* | | -3.838e-07 | | 3.409e-06 | | -0.113 | | 0.910 | |
| *pair_st_laying_date_mod* | | 1.124e-03 | | 8.419e-03 | | 0.133 | | 0.894 | |
| *pair_mean_distance_feeders_preR* | | 4.120e-04 | | 9.449e-04 | | 0.436 | | 0.663 | |

| C. GLM_1st-Mean Weight_ | | | | | | | | | |
| --- | --- | --- | --- | --- | --- | --- | --- | --- | --- |
| Model no. | Explanatory Variables | K | ΔAIC | | df (obs. Deleted) | | Null Dev. | | AIC |
| Dependent variable: *1^st^_brood_weight* | | | | | | | | | |
| 1 | *null* | 0 | - | | 24 (9) | | 23.45 | | 73.35 |
| 2 | *pair_time_preR (ns) +* ***pair_mean_distance_feeders_preR*** *(.)* | 2 | 47.74 | | 10 (23) | | 5.466 | | 25.61 |
| Model summary | | | | | | | | | |
|  | | Estimate | | Std. error | | *t* value | | *P-*value | |
| Intercept | | 1.732e+01 | | 6.105e-01 | | 28.377 | | **2.57e-09 *** | |
| *pair_time_preR* | | -3.878e-06 | | 5.434e-06 | | -0.714 | | 0.4957 | |
| *pair_mean_distance_feeders_preR* | | 2.899e-03 | | 1.520e-03 | | 1.908 | | 0.0929 . | |

| D. GLM_Total Eggs_ | | | | | | | | | |
| --- | --- | --- | --- | --- | --- | --- | --- | --- | --- |
| Model no. | Explanatory Variables | K | ΔAIC | | df (obs. Deleted) | | Null Dev. | | AIC |
| Dependent variable: *pair_egg_tot* | | | | | | | | | |
| 1 | *null* | 0 | - | | 33 (0) | | 67.81 | | 207.6 |
| 2 | ***pair_time_tot*** *(.) +* ***pair_st_laying_date_mod*** *(*) + pair_mean_distance_feeders_tot* | 3 | 110.52 | | 15 (18) | | 35.84 | | 97.08 |
| Model summary | | | | | | | | | |
|  | | Estimate | | Std. error | | *t* value | | *P-*value | |
| Intercept | | 4.619e+00 | | 8.182e-01 | | 5.645 | | **1.65e-08 *** | |
| *pair_time_tot* | | -1.953e-06 | | 1.311e-06 | | -1.489 | | 0.13641 | |
| *pair_st_laying_date_mod* | | -2.069e-02 | | 6.871e-03 | | -3.012 | | **0.00408 *** | |
| *pair_mean_distance_feeders_tot* | | 1.614e-04 | | 6.508e-04 | | 0.248 | | 0.88334 | |
| *pair_year2022* | | 2.662e-01 | | 2.008e-01 | | 1.325 | | 0.18505 | |

| E. GLM_Total Clutches_ | | | | | | | | | |
| --- | --- | --- | --- | --- | --- | --- | --- | --- | --- |
| Model no. | Explanatory Variables | K | ΔAIC | | df (obs. Deleted) | | Null Dev. | | AIC |
| Dependent variable: *pair_clutch_tot* | | | | | | | | | |
| 1 | *null* | 0 | - | | 33 (0) | | 11.55 | | 96.37 |
| 2 | *time_female_tot (ns) + pair_st_laying_date_mod (ns) + pair_mean_distance_feeders_tot (ns)* | 3 | 41.43 | | 15 (18) | | 6.314 | | 54.94 |
| Model summary | | | | | | | | | |
|  | | Estimate | | Std. error | | *t* value | | *P-*value | |
| Intercept | | 2.402e+00 | | 1.829e+00 | | 1.313 | | 0.189 | |
| *time_female_tot* | | -2.119e-06 | | 4.266e-06 | | -0.415 | | 0.678 | |
| *pair_st_laying_date_mod* | | -1.663e-02 | | 1.626e-02 | | -1.023 | | 0.306 | |
| *pair_mean_distance_feeders_tot* | | -4.275e-04 | | 1.595e-03 | | -0.268 | | 0.789 | |
| *year2022* | | 3.432e-01 | | 4.734e-01 | | 0.725 | | 0.469 | |

| E. GLM_1st-2nd_ | | | | | | | | | |
| --- | --- | --- | --- | --- | --- | --- | --- | --- | --- |
| Model no. | Explanatory Variables | K | ΔAIC | | df (obs. Deleted) | | Null Dev. | | AIC |
| Dependent variable: *days_btw_st_nd_laying_dates* | | | | | | | | | |
| 1 | *null* | 0 | - | | 19 (14) | | 183 | | 105 |
| 2 | *time_female_preR (ns) + time_female_R (ns) + time_male_preR (ns) + time_male_R (ns)* | 4 | - 6 | | 19 (14) | | 183 | | 111 |
| Model summary | | | | | | | | | |
|  | | Estimate | | Std. error | | *t* value | | *P-*value | |
| Intercept | | 3.765e+01 | | 1.603e+00 | | 23.494 | | **3.03e-13 *** | |
| *time_female_preR* | | -3.648e-05 | | 5.380e-05 | | -0.678 | | 0.508 | |
| *time_female_R* | | -5.149e-06 | | 2.479e-05 | | -0.208 | | 0.838 | |
| *time_male_preR* | | -1.455e-05 | | 2.815e-05 | | -0.517 | | 0.613 | |
| *time_male_R* | | -1.814e-05 | | 3.674e-05 | | -0.494 | | 0.629 | |
